# *De novo* transcriptome assembly of the Italian white truffle (*Tuber magnatum* Pico)

**DOI:** 10.1101/461483

**Authors:** Federico Vita, Amedeo Alpi, Edoardo Bertolini

## Abstract

The Italian white truffle (*Tuber magnatum* Pico) is a gastronomic delicacy that dominates the worldwide truffle market. Despite its importance, the genomic resources currently available for this species are still limited. Here we present the first *de novo* transcriptome assembly of *T. magnatum*. Illumina RNA-seq data were assembled using a single-*k*-mer approach into 22,932 transcripts with N50 of 1,524 bp. Our approach allowed to predict and annotate 12,367 putative protein coding sequences, reunited in 6,723 loci. In addition, we identified 2,581 gene-based SSR markers. This work provides the first publicly available reference transcriptome for genomics and genetic studies providing insight into the molecular mechanisms underlying the biology of this important species.

## 1. Introduction

*Tuber magnatum* Pico is an edible ectomycorrhizal ascomycete that forms specialized symbioses with higher plants (Bonito and Smith 2016). Also known as Italian white truffle, *Tuber magnatum* is greatly appreciated for its intense flavor and aroma (Splivallo et al. 2011) and among the European truffle species (*Tuber melanosporum*, *Tuber aestivum*, *Tuber borchii*) it is considered the most precious with high economic value (Mello et al. 2006). The white truffle is present only in some parts of Italy, Istria and western Balkans. This species grows in a restricted number of areas and preferably in moist and drained environments with soil characterized by neutral to alkaline pH (Bragato and Marjanović 2016). While the cultivation of the European truffles gave successful results, the Italian white truffle grows only in natural environment. This inability to cultivation together with the high worldwide demand make this truffle one the most expensive food on the market.

To date, *T. melanosporum* and *T. borchii* nuclear genomes have been published (Martin et al. 2010; Murat et al. 2018) revealing crucial information regarding their genome composition, size and organization. Although several ongoing genomics projects will release in the coming years the genome sequences of fourteen additional *Tuber* species, including *Tuber magnatum* Pico (Murat and Martin 2016), at the present time little is known about the evolution, biology and ecology of the *Tuber* genus.

In this frame, this study provides the first publicly available *de novo* transcriptome assembly of *T. magnatum* based on Illumina short-read sequencing giving a valuable resource for future genetic and genomic studies. Moreover, this resource will be useful for providing a genomic reference for gene expression studies in order to investigate several molecular mechanisms such as the aroma biosynthesis pathways and the mechanisms underlying evolution and adaptation of the different Italian white truffle ecotypes.

## 2. Results and Discussion

### 2.1 Illumina Sequencing and Transcriptome Assembly

We built the *T. magnatum* transcriptome generating a cDNA library from fruiting bodies collected in southern, central and northern Italy. To maximize the discovery and accurate estimate of transcripts, all the selected fruiting bodies (see Methods) were homogenized together and subjected to RNA extraction, Illumina poly(A)^+^ library preparation and sequencing. A total number of 375 million 100-bp single-end reads were generated corresponding to ∼ 37 Gbp of high quality data (Phred score > Q30).

We first mapped our sequencing data to *Tuber melanosporum* genome sequence (version 1) (Martin et al. 2010) (downloaded from https://www.genoscope.cns.fr/externe/Download/Projets/Tuber/assembly/) and to *Tuber borchii* (draft) (Murat et al. 2018) (downloaded from http://genome.jgi.doe.gov/Tubbor1) to check the possibility of using the closest species to help the transcriptome reconstruction based on a genome-guided approach. This analysis was conducted using the aligner Hisat2 (Pertea et al. 2016) with default parameters (maximum number of mismatches allowed equal to 2) and resulted in 2.24% and 1.36% of overall alignment rate against *T. melanosporum* and *T. borchii*, respectively (Table S1). Thus, we proceeded with a *de novo* transcriptome assembly using a single kmer (k=25) employing the program Trinity (Haas et al. 2013). Overall, 99,507 transcripts were pre-assembled and, in order to eliminate the transcriptome redundancy and the artifacts derived from sequencing together three *T. magnatum* ecotypes, the longest transcript for each locus was selected according to 95% sequence identity. The remaining sequences were then screened through the EvidentialGene pipeline in order to generate the best biological dataset comprising both coding and non-coding transcripts, leading to 67,120 unique transcripts (unitranscripts). Candidate coding regions were predicted using the tool TransDecoder for a total of 22,932 transcripts with N50 of 1,524 bp and a mean transcript length of 696 bp (Table 1 and Figure 2). This set was then subsetted for sequences containing complete ORFs resulting in 12,367 transcripts, reunited 6,723 loci (see deposited files in the figshare db).

**Table 1.** *Tuber magnatum de novo* transcriptome assembly statistics

| **Metric** | **Final transcriptome** |
| --- | --- |
| Total number of transcripts | 22,932 |
| Total assembled bases (bp) | 24,090,894 |
| Max transcript length (bp) | 17,115 |
| Mean transcript length (bp) | 696 |
| N50 (bp) | 1,524 |
| Total number of cds | 18,197 |
| Total number of cds < 5 kb | 17,977 |
| Total number of high-confidence genes* | 6,723 |
| Total number of high-confidence transcripts* | 12,367 |

\*sequences containing a complete ORF

Overall, the number of gene models we annotated in*T. magnatum* is comparable with the closely related species *T. melanosporum* (version 1, 7,496 gene models annotated) whilst seems drastically reduced compared to *T. borchii* (draft, 12,346 gene models annotated) (Murat et al. 2018). To evaluate the consistency of the assembly we first mapped all the RNA-seq reads back to the transcriptome (22,932 transcripts) using the algorithms Salmon and Bowtie2. The alignment rate for the programs was 68.35% and 61.62%, respectively with 51.61% of unique mapped reads resulted from the latter one. Overall, considering we mapped back the reads only on a small fraction of the transcriptome (34% of total unitranscripts), this result confirms the high quality of the *T. magnatum*sequence transcripts and their assembly.

We also checked the transcriptome completeness and gene prediction by comparing our set of transcripts with the set of genes belonging to Ascomycota and Pezizomycotina by running the Benchmarking Universal Single-Copy Orthologs tool (BUSCO) (Simão et al. 2015; Waterhouse et al. 2017). Out of a total of 4,471, the number of complete and single copy BUSCOs found in our dataset was 3,217 (72%) while the number of missing BUSCOs was 689 (15%) with ∼ 11% of fragmented BUSCOs found.

In addition, the putative protein coding sequences (12,367) were functionally annotated using Trinotate (Bryant et al. 2017) searching for nucleotide and protein sequence homology against the UniProtKb/Swiss-Prot and Pfam database. When available, Gene Ontology (GO) (see Supplemental Figure S1) and KEGG terms were retrieved from TrEMBL/SwissProt database and associated to the transcripts (see deposited files in the figshare db).

### 2.2 Genic-single sequence repeat (SSR)

To date, no genetic markers have been reported for *Tuber magnatum* Pico. We exploited our transcriptome data for the development and characterization of gene-based SSR markers. The candidate coding transcripts (22,932) were screened for repeat motifs to explore the microsatellite profile in *T. magnatum*. A total of 2,581 SSRs were obtained from 2,225 candidate coding transcripts (∼ 10%). Our data suggest strong enrichment of mono-and tri-nucleotide repeat motifs among the five SSR classes we analyzed (see Method), while longer repeat motifs were underrepresented (see deposited files in the figshare db).

### 2.3 Data

Raw RNA-seq data generated in this study were deposited in the NCBI Sequence Read Archive (SRA) (BioProject ID: PRJNA501857). All the transcriptome data developed in this study were deposited in the figshare database and are accessible through the link https://figshare.com/s/01ec86ee5e0f657ab955

## 3. Methods

### 3.1 Biological Material

*T. magnatum* fruiting bodies were collected in southern, central and northern Italy, respectively in the regions of Molise, Tuscany and Piedmont in the years 2013, 2014 and 2015. According to the percentage of ascidia as described by (Zeppa et al. 2002) we selected 6 carpophores at stage five (average weight 7-10 g) for each year, for a total of 18 carpophores per location. Fruiting bodies were thoroughly washed with distilled water and subsequently rinsed with absolute ethyl alcohol to remove contaminants. The thin external layer of the peridium was removed and samples were flash frozen in liquid nitrogen and stored at −80 °C until further processing.

From each fruiting body we collected 100 mg of tissue and pooled them together by year and geographical area for a total of 9 representative samples.

### 3.2 RNA Extraction, Library Construction and Sequencing

The nine samples were homogenized with liquid nitrogen using mortar and pestle. To minimize the variability derived from the sampling time and locations, the samples were balanced and mixed. RNA extraction was achieved using the Plant/Fungi Total RNA Purification Kit (Norgen Biotek Corp., Canada) according to the manufacturer protocol. Total RNA integrity was further checked on 1% agarose gel and using Agilent Bioanalyzer 2100 (Agilent Technologies, Santa Clara, CA). An Illumina directional poly(A)^+^ RNA library was generated from the mixed sample according to the TruSeq RNA Library Prep Kit (Illumina, San Diego, CA) and subjected to sequencing using a single-read 100 bp design with a HiSeq2000 sequencer at IGATech (Udine, Italy). Illumina CASAVA v1.8.2 pipeline was used to process raw data for format conversion.

### 3.3 Read Pre-Processing and *de novo* Assembly

Raw reads were quality evaluated before the data analysis using the program FastQC v0.11.5 (Andrews et al. 2014). A quality score above Q30 was kept to maintain high accuracy in the downstream analysis. Undefined bases (Ns) within the reads and the presence of sequencing adapters were excluded with the program Cutadapt (version 1.8.3) (Martin 2011).

All the quality reads were employed to *de novo* reconstruct the *Tuber magnatum* transcriptome with a de Bruijn graph based approach. Trinity assembler (v2.1.1) (Haas et al. 2013) with a single *k-mers*method (*k-mer* = 25) was used. The resulting assembled transcripts were then clustered based on 95% sequence identity using the program CD-HIT (v4.6.6) (Fu et al. 2012). To generate a final assembly, the reconstructed sequences were further evaluated through the EvidentialGene tr2aacds pipeline (http://eugenes.org/EvidentialGene/) leading to a final transcriptome assembly containing a unique set of transcripts (unitranscripts). Potential protein coding transcripts were predicted using the *ab initio* utility TransDecoder (https://github.com/TransDecoder) to extract long open reading frames (ORFs) and predicts coding regions.

Trinotate (v.3) (Bryant et al. 2017) was applied to functional annotate the non-redundant *T. magnatum* transcriptome searching against the UniProtKb/Swiss-Prot database (uniprot_sprot.pep, downloaded from: https://data.broadinstitute.org/Trinity/Trinotate_v3_RESOURCES/) using the algorithim BLASTx and BLASTp (v2.7.1+) with the options: −evalue 1e10 −num_threads 4 −max_target_seqs 1 −outfmt 6. Functional domains were searched using HMMER (v3.2.1) (Finn et al. 2015) against the Pfam datatabase (Pfam-A.hmm, downloaded from: https://data.broadinstitute.org/Trinity/Trinotate_v3_RESOURCES/) with default options. All the top-matching BLAST hits and the corresponding Gene Ontology (GO) and Kyoto Encyclopedia of Genes and Genomes (KEGG) assignments from TrEMBL/SwissProt databases were extracted using a maximum e-value cutoff of 1e-5.

Transcript abundance and quality of the assembled transcriptome were estimated using salmon (v0.9.1) (Patro et al. 2017) in *quasi-mapping* mode with the option ––numBootstraps 30 and Bowtie2 (Langmead and Salzberg 2012) in end-to-end mode (––sensitive) with maximum number of mismatches allowed equal to 1.

The quality of the non-redundant *T. magnatum* transcriptome was checked using the Benchmarking Universal Single-Copy Orthologs tool (BUSCO) (v3.0.2) (Simão et al. 2015; Waterhouse et al. 2017) comparing the unitranscripts with the Ascomycota and Pezizomycotina specific lineage conserved single copy orthologs derived from OrthoDB v9.

### 3.4 Identification of genic-SSRs

*T. magnatum* transcripts were scanned for single sequence repeats (SSRs) markers using the program MISA (Beier et al. 2017) with default options searching for mono-, di-, tri-, tetra-, penta-, hexa-nucleotide motifs with minimum number of repeat units equal to 10, 6, 5, 5,5,5 respectively.

## Author Contributions

FV designed the experiment and performed the wet-lab activities. EB designed the experiment, performed the *in silico* analyses and wrote the manuscript. AP read critically and drafted the paper.

## Acknowledgements

Sequencing cost was supported by Tuscany regional administration through the research project VOLATOSCA (http://www.ipsp.cnr.it/projects/volatosca). Data analysis was conducted and supported by the International Doctoral Programme in Agrobiodiversity – Scuola Superiore Sant’Anna (www.santannapisa.it) and LOLIETTOO ^®^ – the academic spin-off company of Scuola Superiore Sant’Anna (www.loliettoo.it).

---

**Figure 1.**
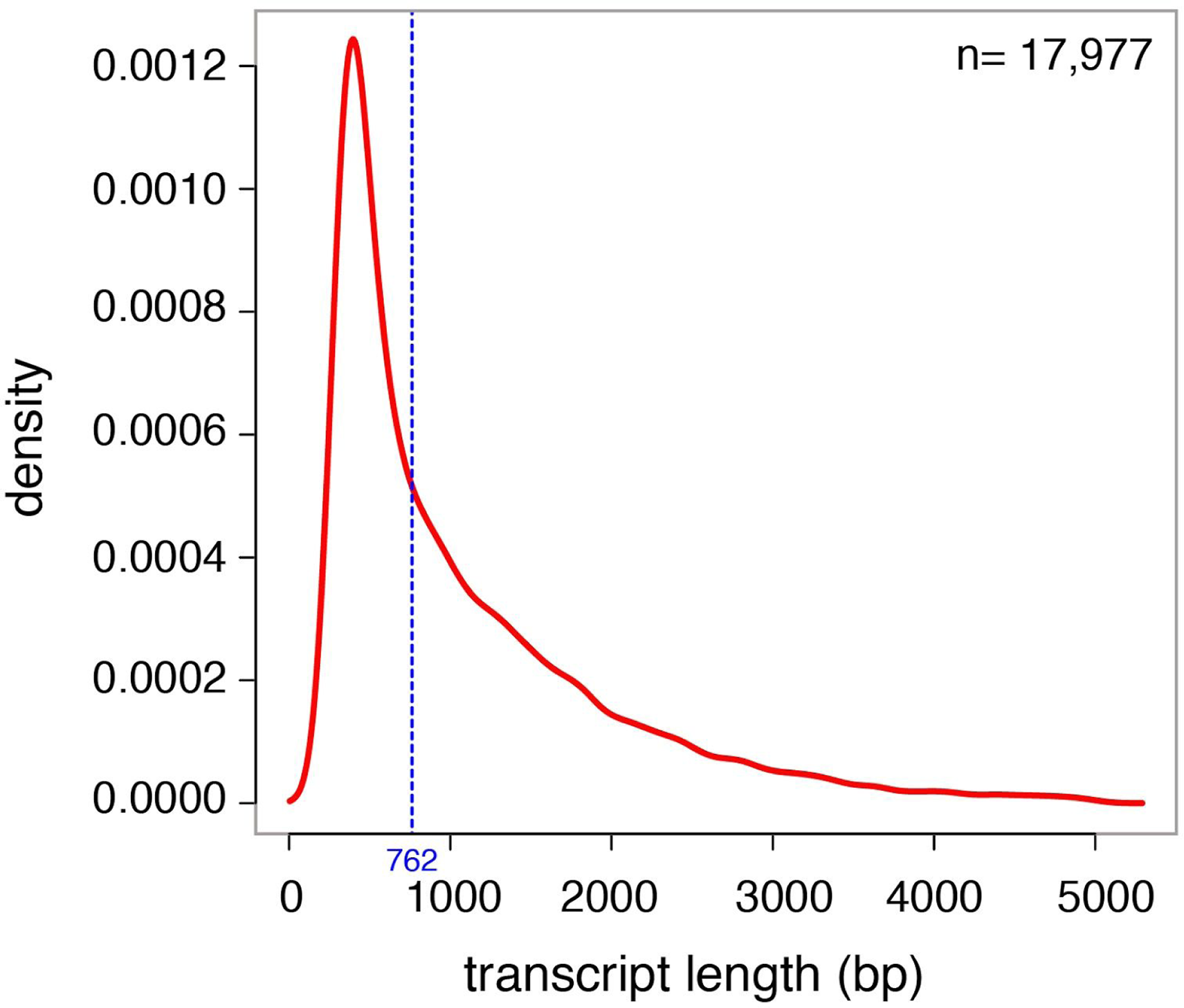
Transcript length distribution of *Tuber magnatum de novo* transcriptome. The image represents the cds transcripts (n=17,988) selected for sequence length < 5000 bp (98% of cds). Blue dotted line represents the median length.

**Figure S1.**
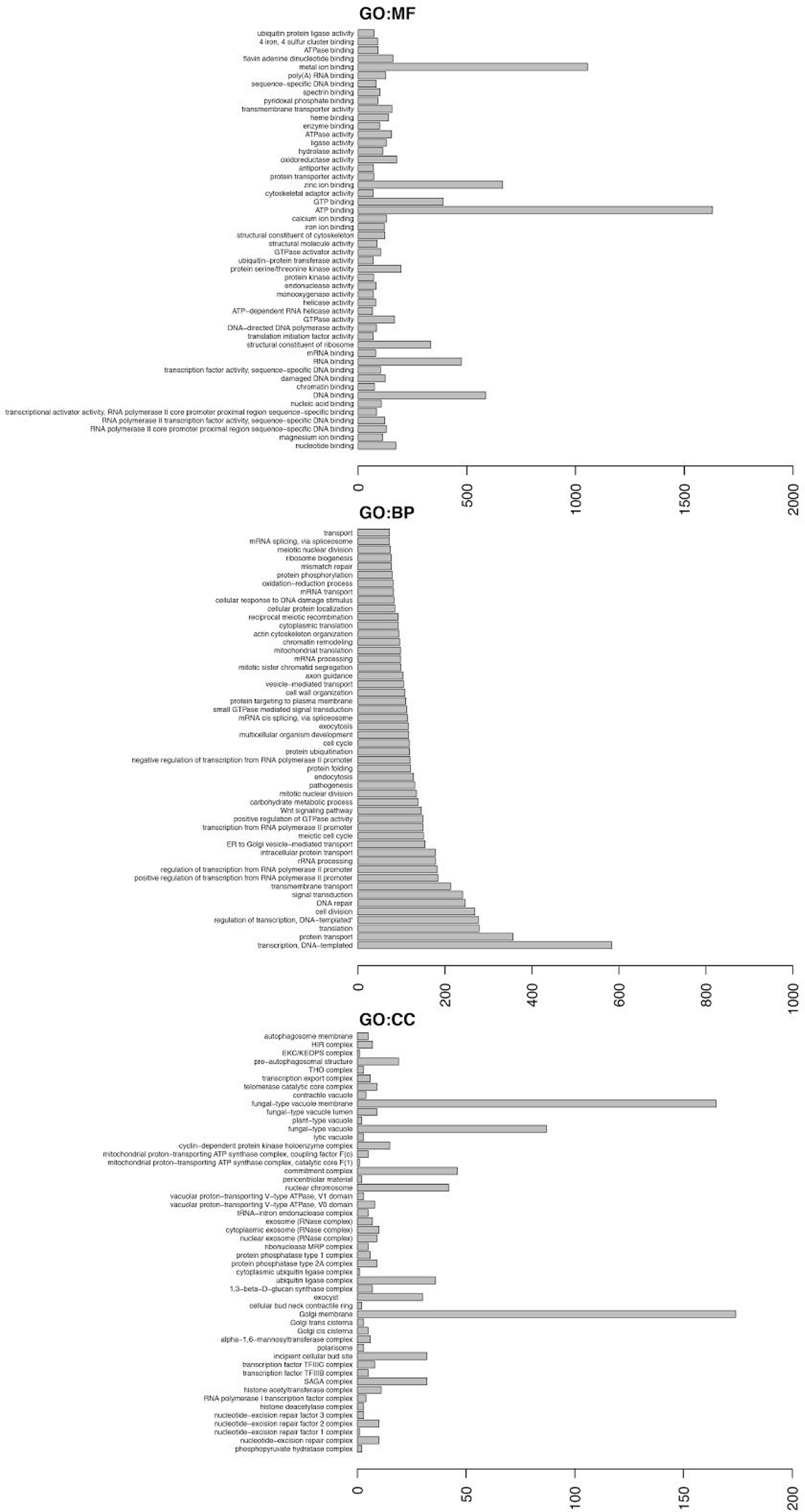
Histograms represent the frequency of the top 50 GO terms associated with the annotated *T. magnatum* transcripts. Gene Ontology categories: molecular functions (MF), biological process (BP), cellular components (CC) are represented. X-axis represents the frequency of the GO term. Y-axis shows the description of the GO term.

**Table S1.** Mapping statistics of *T. magnatum* against the reference genomes of *T. borchii* and *T. melanosporum*. See Result section for details.

|  | **Total**<br>**number of**<br>**reads** | **Reads mapped**<br>**exactly 1 time** | **Reads**<br>**mapped >1**<br>**times** | **Unmapped**<br>**reads** | **Overall**<br>**alignment rate** |
| --- | --- | --- | --- | --- | --- |
| *T. borchii* | 375,046,769 | 5,075,954 | 41,206 | 369,929,609 | 1.36% |
| *T. melanosporum* | 375,046,769 | 6,846,901 | 1,551,868 | 366,648,000 | 2.24% |

